# Working Towards a Blood-Derived Gene Expression Biomarker Specific for Alzheimer’s Disease

**DOI:** 10.1101/621987

**Authors:** Hamel Patel, Raquel Iniesta, Daniel Stahl, Richard J.B Dobson, Stephen J Newhouse

**Affiliations:** Department of Biostatistics and Health Informatics, Institute of Psychiatry, Psychology and Neuroscience, King’s College London, London, UK; NIHR BioResource Centre Maudsley, NIHR Maudsley Biomedical Research Centre (BRC) at South London and Maudsley NHS Foundation Trust (SLaM) & Institute of Psychiatry, Psychology and Neuroscience (IoPPN), King’s College London; Health Data Research UK London, University College London, 222 Euston Road, London, UK; Institute of Health Informatics, University College London, 222 Euston Road, London, UK; The National Institute for Health Research University College London Hospitals Biomedical Research Centre, University College London, 222 Euston Road, London, UK

## Abstract

**Background:** A significant number of studies have investigated the use of blood-derived gene expression profiling as a biomarker for Alzheimer’s Disease (AD). However, the typical approach of developing classification models trained on subjects with AD and complimentary cognitive healthy controls may result in markers of general illness rather than being AD-specific. Incorporating additional related neurological and age-related disorders during the classification model development process may lead to the discovery of an AD-specific expression signature.

**Methods:** Two XGBoost classification models were developed and optimised. The first used the typical approach, training on 160 AD and 160 cognitively normal controls, while the second was trained in 6318 AD and 6318 mixed controls. Up-sampling was performed in each training set to the minority classes to avoid sampling bias, and both classification models were evaluated in an independent dataset consisting of 127 AD and 687 mixed controls. The mixed control group represents a heterogeneous ageing population consisting of Parkinson’s Disease, Multiple Sclerosis, Amyotrophic Lateral Sclerosis, Bipolar Disorder, Schizophrenia, Coronary Artery Disease, Rheumatoid Arthritis, Chronic Obstructive Pulmonary Disease, and cognitively healthy subjects.

**Results:** The typical approach resulted in a 74 gene classification model with a validation performance of 58.3% sensitivity, 30.3% specificity, 13.4% PPV and 79.7% NPV. In contrast, the second approach resulted in a 28 gene classification model with an overall improved validation performance of 46.5% sensitivity, 95.6% specificity, 66.3% PPV and 90.6% NPV.

**Conclusions:** The addition of related neurological and age-related disorders into the AD classification model developmental process identified a more AD-specific expression signature, with improved ability to distinguish AD from other related diseases and cognitively healthy controls. However, this was at the cost of sensitivity. Further improvement is still required to identify a robust blood transcriptomic signature specific to AD.

## Introduction

Alzheimer’s disease (AD) is a progressive neurodegenerative disorder affecting an estimated one in nine people over the age of 65 years of age, making it the most common form of dementia worldwide [1]. Current clinical diagnosis of the disease is primarily based on a time-consuming combination of physical, mental and neuropsychological examinations. With the rapid increase in the prevalence of disease, there is a growing need for a more accessible, cost-effective and time-effective approach for early diagnosis and monitoring AD.

For research purposes, brain positron emission tomography (PET) scans and cerebral spinal fluid (CSF) have been used for disease identification. In particular, decreased Aβ and increased tau levels in CSF have been successfully used to distinguishing between AD, mild cognitive impairment (MCI) and cognitive healthy individuals with high accuracy. However, as a relatively invasive and costly procedure, it may not appeal to the majority of patients or be practical on a large-scale trial basis for screening the population [2] [3] [4]. Peripheral blood-derived biomarkers could potentially be a solution to this problem.

Blood is a complex mixture of fluid and multiple cellular compartments that are consistently changing in protein, lipid, RNA and other biochemical entity concentrations [5], which may be useful for AD diagnosis. A recent study reviewed 163 candidate blood-derived proteins as potential AD biomarkers from 21 separate studies [6]. The overlap of biomarkers between studies was limited, with only four biomarkers α-1-antitrypsin, α-2-macroglobulin, apolipoprotein E and complement C3 found to replicate in five independent cohorts. However, a follow-on study discovered these biomarkers were not specific to AD, and were also discovered to be associated with other brain disorders including Parkinson’s Disease (PD) and Schizophrenia (SCZ) [7], suggesting the need to consider other neurological and related disorders in study designs to enable the discovery of biomarkers specific to AD.

Similarly, several studies have attempted to exploit blood transcriptomic measurements for AD biomarker discovery. Initial research was limited to the analysis of single differentially expressed genes (DEG) as a means to distinguish AD from cognitively healthy individuals [2] [8]. However, the limited overlap and reproducibility of DEG from independent cohorts suggests this method alone is not reliable enough [2]. A solution to this problem would be to use information across all genes simultaneously through machine learning algorithms to identify combinations of gene expression changes that may represent a biomarker for AD. This technique has been employed in multiple studies, which have demonstrated the ability to differentiate AD from non-AD subjects [3] [9] [10] [11] [12] [3] [9]. However, small sample size and lack of independent validation datasets most likely led to overfitting. The decrease in costs associated with microarray technologies led a study developing an AD classification model based on a larger training set of 110 AD and 107 controls and validating in a larger independent cohort of 118 AD and 118 controls. The model achieved 56% sensitivity, 74.6% specificity, and an accuracy of 66%, which equated to 69.1% Positive Predictive Power (PPV) and 63% Negative Predictive Power (NPV) [10]. This was one of the first studies to demonstrate validation in an independent cohort; however, the classification model still lacked the 90% predictive power desired from a clinical diagnostic test [13].

Previous studies have demonstrated the potential use of blood transcriptomic levels to differentiate between AD and cognitively healthy individuals; however, they are yet to be precise enough for clinical utility and are yet to be extensively evaluated on specificity by assessing model performance in a heterogeneous ageing population of multiple diseases. This validation process is critical to determine whether the classification model is indeed disease-specific, a general indication of ill health, or an overfit.

This study developed a novel XGBoost classification model trained on blood transcriptomic profiling from AD, related mental disorders (Parkinson’s disease [PD], Multiple Sclerosis [MS], Amyotrophic Lateral Sclerosis [ALS], Bipolar Disorder [BD], Schizophrenia [SCZ]), age-related disorders (Coronary Artery Disease [CD], Rheumatoid Arthritis [RA], Chronic Obstructive Pulmonary Disease [COPD]), and cognitively healthy subjects to differentiate AD from diseased and otherwise normal subjects. The classification model was developed with clinical utility in mind, with each dataset processed and transformed independently and evaluated in an independent ageing heterogenous population (testing set) consisting of similar diseases as the training set.

## Methods

### Data acquisition

Microarray gene expression studies were sourced from publicly available repositories GEO (https://www.ncbi.nlm.nih.gov/geo/) and ArrayExpress (https://www.ebi.ac.uk/arrayexpress/) in May 2018. Study inclusion criteria were; 1) microarray gene expression profiling must be performed on an age-related or neurological disorder, 2) RNA was extracted from whole blood or a component of blood, 3) study must contain at least ten subjects, and 4) data was generated on either the Illumina or Affymetrix microarray platform using a BeadArray containing at least 20,000 probes. The microarray platform was restricted to Affymetrix and Illumina only, as replication between the two platforms is generally very high [14], and BeadArrays restricted to a minimum of 20,000 probes to maximise the overlap of genes across studies, while also optimising the number studies available for inclusion.

### Data processing

The data processing pipeline was designed with reproducibility and clinical utility in mind. New subjects could be independently processed and predicted through the same classification model without using any prior knowledge on gene expression variation of the data used to develop the classification model and without making any alteration to the classification model itself. All data processing was undertaken in RStudio (version 1.1.447) using R (version 3.4.4). Microarray gene expression studies were acquired from public repositories using the R packages “GEOquery” (version 2.46.15) and “ArrayExpress” (version 1.38.0). For longitudinal studies involving treatment effect, placebo subjects or initial gene expression profiling from baseline subjects before treatment were used. Studies consisting of multiple disorders were separated by disease into datasets consisting of diseased subjects and corresponding healthy controls if available.

Raw gene expression data generated on the Affymetrix platform were “mas5” background corrected using R package “affy” (version 1.42.3), log2 transformed and then Robust Spline Normalised (RSN) using R package “lumi” (version 2.16.0). Datasets generated on the Illumina platform were available in either a “raw format” containing summary probes and control intensities with corresponding p-values or a “processed format” where data had already been processed and consisted of a subset of probes and samples deemed suitable by corresponding study authors. When acquiring studies, preference was given to “raw format” data where possible, and when available, was “normexp” background corrected, log2 transformed, and quantile normalised using the “limma” R package (version 3.20.9).

Sex was then predicted using the R package “massiR” (version 1.0.1) and subjects with discrepancies between predicted and recorded sex removed from further analysis. Next, within each gender and disease diagnosis group of a dataset, probes above the “X” percentile of the log2 expression scale in over 80% of the samples were deemed “reliably detected”. To account for the variation of redundant probes across different BeadArrays, the “X” percentile threshold value was manually adjusted until a variety of robust literature defined house-keeping genes were correctly defined as expressed or unexpressed in their corresponding gender groups [15]. Any probe labelled as “reliably detected” in any group (based on gender and diagnosis) was taken forward for further analysis from all samples within that dataset. This substantially eliminates noise [16] and ensures disease and gender-specific signatures are captured within each dataset.

Next, to ensure homogeneity within biological groups, outlying samples were iteratively identified and removed using the fundamental network concepts described in [17]. Finally, to enable cross-platform probes to be comparable, platform-specific probe identifiers were annotated to their corresponding universal Entrez gene identifiers using the appropriate BeadArray R annotation files; “hgu133plus2.db”, “hgu133a.db”, “hugene10sttranscriptcluster.db”, “illuminaHumanv4.db” and “illuminaHumanv3.db”.

### Cross-platform normalisation

To enable transcriptomic information between datasets to be directly comparable, a rescaling technique, the YuGene transform, was applied to each dataset independently. YuGene assigns modified cumulative proportion value to each measurement, without losing essential underlying information on data distributions and allows transformation of independent studies and individual samples [18]. This allows new data to be added without global renormalisation and enables the training and testing data to be independently rescaled. Common probes across all processed datasets that contained both female and male subjects were extracted from each dataset and independently rescaled using the R package YuGene (version 1.1.5). YuGene transformation assigns a value between 0 and 1 to each gene, where 1 is highly expressed. As samples originated from publicly available datasets, potential duplicate samples may exist in this study. To address this issue, correlation analysis was performed on all samples using the common probes. Any sample with a Pearson’s correlation coefficient equal to 1 was suggested to be a duplicate sample and would be removed from further analysis.

### Training Set and Testing Set assignment

Multiple datasets from the same disease were available, allowing entire datasets to be assigned to either the “Training Set” for classification model development or “Testing Set” for independent external validation purposes. Larger datasets from the same disease were prioritised to the training set, allowing the machine learning algorithm to learn in a larger discovery set.

Individual subjects within the training and testing set were assigned a “0” class if the individual was AD or “1” if the individual was non-AD (includes healthy controls and non-AD diseased subjects). Grouping the non-AD subjects into a single class effectively mimics a large heterogeneous ageing population where subjects may have a related mental disorder, neurodegenerative disease, age-related disease or are considered relatively healthy.

### Classification model development

Two classification models were created. (i) The first was developed using only the AD and associated control datasets available in the training set and is referred to as the “AD vs healthy control” classification model. (ii) The second classification model was developed using all datasets and diseases available in the training set (AD, non-AD disease and all controls) and is referred to as the “AD vs mixed control” classification model. The second approach aimed to develop a classification model that may be more specific in identifying AD than the typical “AD vs healthy control” classification model.

Classification models were built using a powerful tree boosting algorithm, XGBoost, which in 2015 was used in every winning team in the top 10 of the Data Mining and Knowledge Discovery competition for a wide range of machine learning problems [19] and was suggested to be one of the most sophisticated methods at the time of this work [20], [21]. Furthermore, the tree learning algorithm uses parallel and distributed computing and is approximately 10 times faster than existing methods and allows many hyperparameters to be tuned to reduce the chance of overfitting [20].

First, the training sets were balanced by up-sampling the minority class with replacement to match the number of samples in the majority class. As the second classification model (“AD vs mixed control”) consisted of multiple diseases and complementary controls in the training set, all the control samples were assumed to be healthy and were therefore pooled. Then, within each disease classes of the mixed control group were up-sampled to match the total number of samples in the pooled control group. The AD samples were then up-sampled to match the total number of samples in the mixed control group. This process would ensure that all diseases in the mixed control group had the same probability to be used during the model development process.

Next, the R package “xgboost” (version 0.6.4.1) was used to create optimised models. Default tuning parameters were set to **eta**=0.3, **max_depth**=6, **gamma**=0, **min_child_weight**=1, **subsample**=1, **colsample_bytree**=1, **objective**=“binary: logistic”, **nrounds**=5000, **early_stopping_rounds parameters**=20 and **eval_metric=**“logloss”. The “**seed**” was randomly assigned “222” throughout the model developmental stages for reproducibility purposes. The initial model was built and internally evaluated using 10-fold cross-validation with stratification which calculates a test logloss mean at each **nrounds** iteration, stopping if an improvement to the test logloss means is achieved in the last 20 iterations. The **nrounds** iteration that achieved the optimal test logloss mean was used to build the initial classification model, reducing the chance for an “overfit” model.

During the internal cross-validation process, each feature (gene) was assigned an importance value (“variable importance feature”) based on how well it contributed to the correct prediction of individuals in the training set. The higher the variable importance value for a gene, the more useful that gene was in distinguishing AD subjects from non-AD individuals. The genes contributing to the initial XGBoost model were each assigned a variable importance value. The least two variable important features were then iteratively removed, classification models re-built, and logloss performance measures re-evaluated. This process was repeated through all available baseline features, with the minimum logloss from all iterations used to determine the most predictive genes. This process is referred to as “recursive feature elimination” and has been shown to improve classification model performance and reduce model complexity by removing weak and non-predictive features [22].

Following identification of the most predictive genes, the classification model was further refined by iteratively tuning through the following hyperparameter values: **max_depth** (2:20, 1), **min_child_weight** (1:10, 1), **gamma** (0:10, 1), **subsample** (0.5:1, 0.1), **colsample_bytree** (0.5:1, 0.1), **alpha** (0:1, 0.1),), **lambda** (0:1, 0.1), and **eta** (0.01:0.2, 0.01), whilst performing a 10-fold cross-validation with stratification and evaluating the test logloss mean to select the optimum hyperparameters.

### Classification model evaluation

The classification models were validated on the independent unseen testing set, predicting the diagnosis of all subjects as a probability ranging from 0 to 1, where AD ≤ 0.5 > non-AD. The prediction accuracy, sensitivity, specificity, PPV and NPV were calculated to evaluate the overall classification model’s performance. To assess the sensitivity and specificity of the classifiers, ROC curves and AUC scores were generated using the R package “ROCR” (version 1.07) with the following recommended diagnostic interpretations used: “excellent” (AUC = 0.9-1.0), “very good” (AUC = 0.8-0.9), “good” (AUC = 0.7-0.8), “sufficient” (AUC = 0.6-0.7), “bad” (AUC = 0.5-0.6), and “test not useful” when AUC value is <0.5 [23].

The clinical utility metrics were calculated to evaluate the clinical utility of the classification models. The positive Clinical Utility Index (CUI +) was calculated as PPV * (sensitivity/100) and the negative Clinical Utility Index (CUI -) calculated as NPV * (sensitivity/100). The Clinical Utility Index (CUI) essentially corrects the PPV and NPV values for occurrence of that test in each respective population and scores can be converted into qualitative grades as recommended: “excellent utility” (CUI >= 0.81), “good utility” (CUI >=0.64) and “satisfactory utility” (CUI >=0.49) and “poor utility” (CUI < 0.49) [24]. An overview of the classification model development and evaluation process is provided in Figure 1.

**Figure 1:**
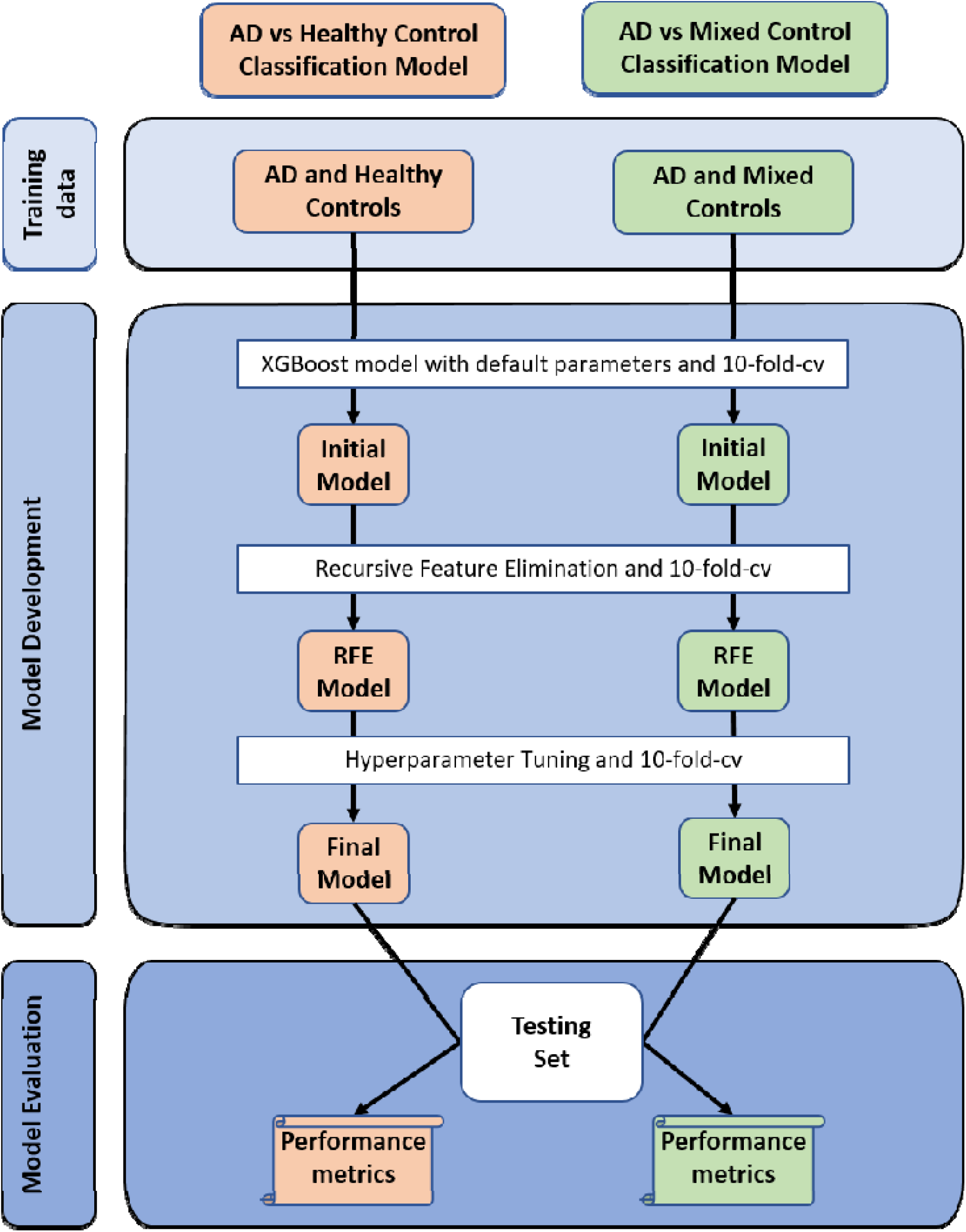
Overview of Study Design. “Logloss” metric was used throughout the model developmental stage to identify optimal features and hyperparameters. Abbreviations: cv = cross-validation and RFE = Recursive Feature Elimination.

### The biological importance of predictive features

The final classification model taken forward for external validation contains a list of ranked genes which collectively differentiate AD from non-AD subjects. These genes have been derived from multiple disorders and may be involved with biological processes, which was therefore assessed though Gene Set Enrichment Analysis (GSEA). The predictive genes were analysed using an Over-Representation Analysis (ORA) implemented through the ConsensusPathDB (http://cpdb.molgen.mpg.de) web-based platform (version 33) [25] in November 2018. For GSEA analysis, a minimum overlap of the query signature and database was set as 2.

## Results

### Summary of data processing

Twenty-one publicly available studies were identified, acquired and processed. Separating studies by disease status resulted in 22 datasets, which consisted of 3 AD, 3 MS, 3 SCZ, 3 CD, 3 RA, 2 COPD, 2 BD, 2 PD and 1 ALS orientated dataset. Only 15 datasets contained both diseased and complementary control subjects, while the remaining 7 contained only diseased subjects. An overview of the demographics of each dataset is illustrated in Table 1.

**Table 1:**
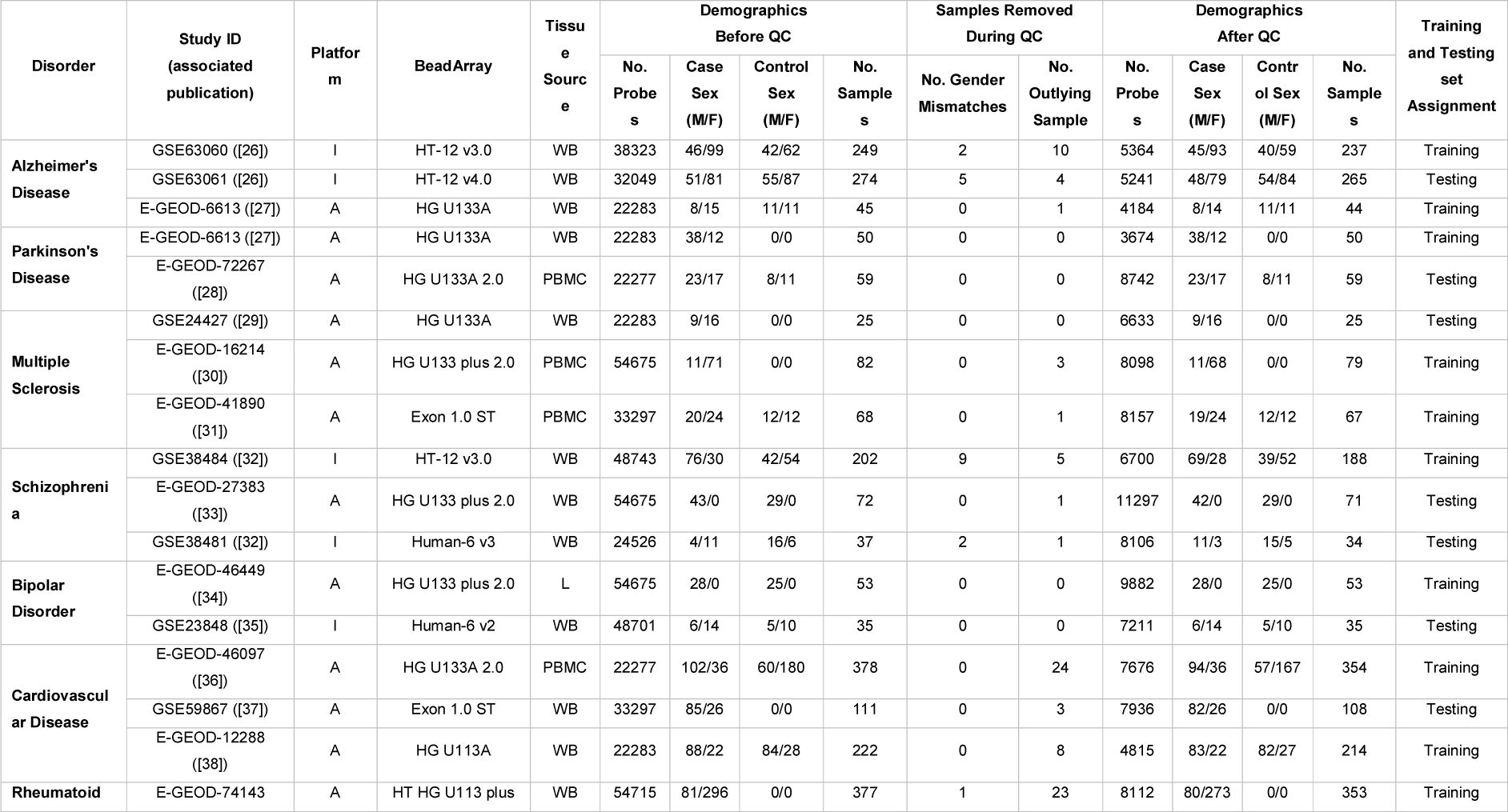
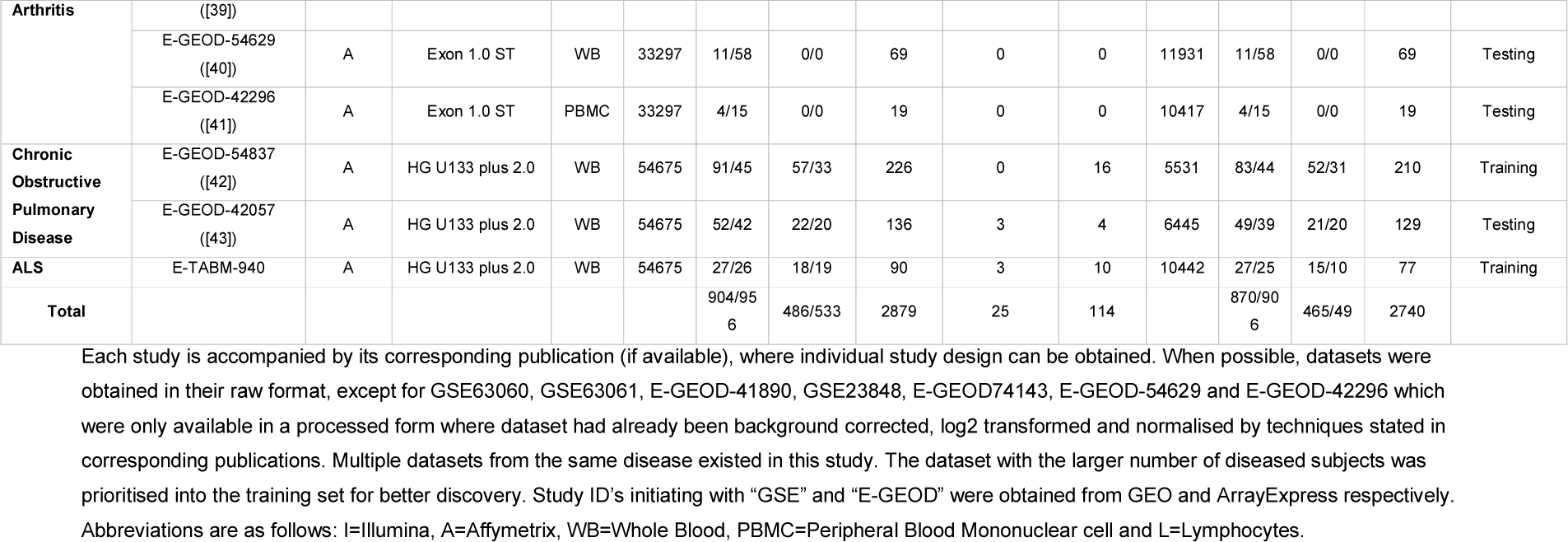
Dataset Demographics

Independently processing the 22 datasets resulted in a total of 2740 samples after Quality Control (QC), of which 287 samples were AD. Since 11 different BeadArrays had been used to expression profile the 9 different diseases, and as 7 datasets were only available in a “processed format” (GSE63060, GSE63061, E-GEOD-41890, GSE23848, E-GEOD74143, E-GEOD-54629 and E-GEOD-42296), each dataset varied in the number of “reliably detected” genes after QC (detailed in Table 1). Initially, an overlap of the common measurable probes across all 22 datasets which were also deemed “reliably detected” in any one of the datasets was compiled, resulting in 7452 genes. In theory, this would ensure all measurable sex and disease-specific genes are captured. However, following independent transformation of each dataset, platform and BeadArray-specific batch effects were observed (**Figure** 2a-b). This can be largely explained by different platforms having different probe designs to target different transcripts of the same gene, leading to significant discrepancies and even absence in the measurement of the same gene by different platforms [14]. Therefore, to address this platform and BeadArray-specific batch effect, 1681 common “reliably detected” genes across all datasets that contained both male and female subjects (20 datasets) were extracted from each dataset and independently YuGene transformed. Essentially, these 1681 genes are expressed at a level deemed “reliably detected” in all 11 different BeadArrays. The distribution of the 1681 genes in each subject is shown in **Figure** 2c-d and can be seen to be more evenly distributed across the 2740 subjects than **Figure** 1a-c, a characteristic desired for the machine learning algorithms. Correlation analysis was then performed on all samples, which suggested all samples were highly correlated, with the maximum per sample correlation coefficients ranging from 0.86-0.99. No sample was deemed to be a duplicate, and therefore, no additional sample was removed following QC.

**Figure 2:**
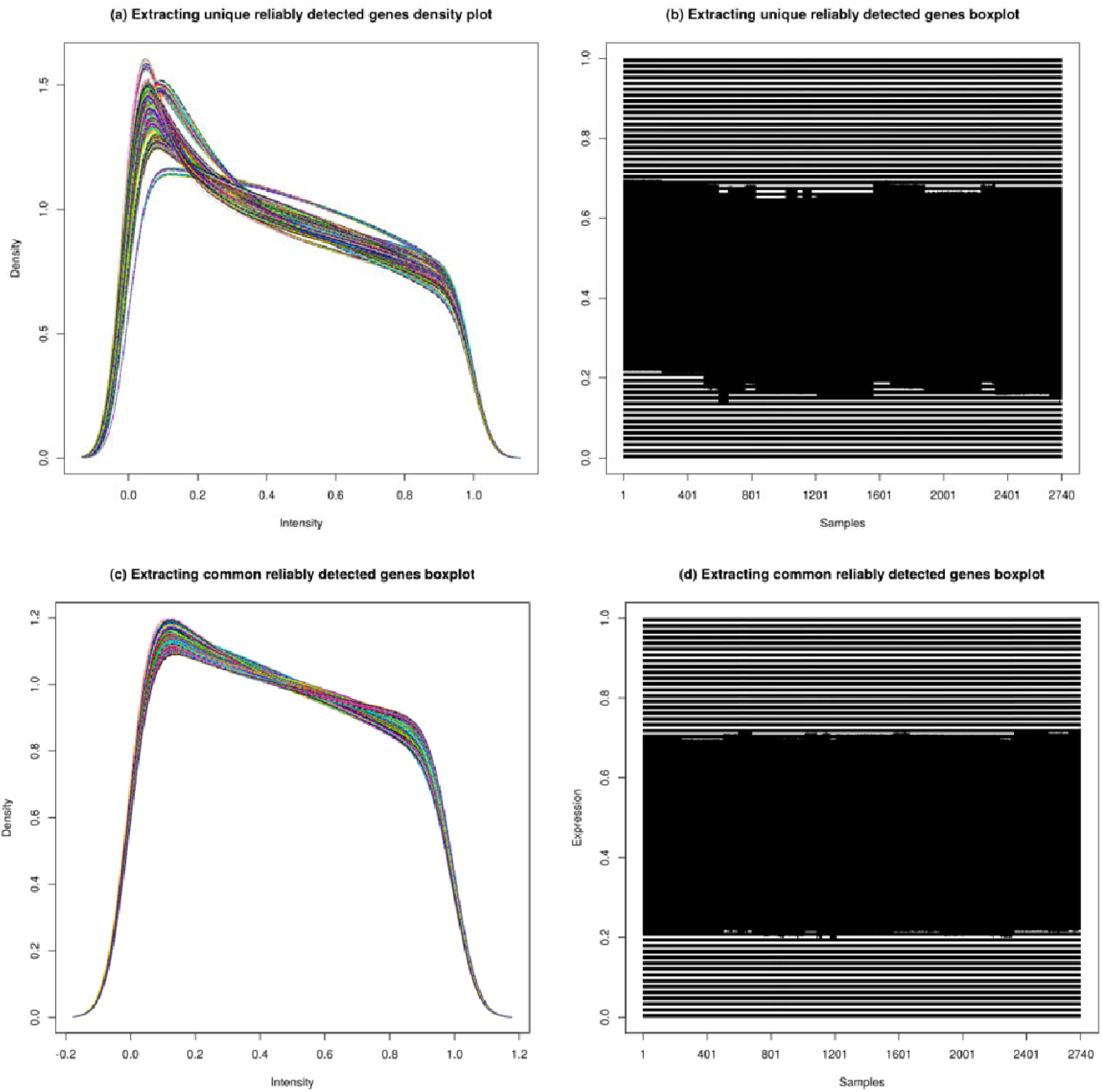
Distribution of gene expression across all subjects. a) and b) demonstrates batch effects caused by specific platform and BeadArrays when extracting 7452 unique genes deemed “reliably detected” in any of the 22 datasets. The shallow “n” curve in density plot a) are dominantly Illumina generated data. In contrast, c) and d) reveals a more evenly distributed gene expression profile across all subjects when extracting the 1681 common “reliably detected” genes, and therefore was used for classification model development.

### Training Set and Testing Set demographics

Multiple datasets from the same disease were obtained for this study, with the largest dataset from each disease assigned to the training set to improve discovery. However, three AD datasets were available, and the two largest datasets were generated on the Illumina platform, and the third on the Affymetrix platform. To address any subtle differences in gene expression which may still exist in the data due to platform differences, the largest Illumina AD and the Affymetrix AD datasets were both assigned to the Training Set.

Following dataset assignment, the training set consisted of 160 AD subjects and 1766 non-AD subjects, while the testing set consisted of 127 AD subjects and 687 Non-AD subjects. The Non-AD group in both the training and testing set consisted of subjects with either PD, MS, SCZ, BD, CD, RA, COPD or were relatively healthy. Only one ALS dataset suitable for this study was identified and was deemed too small to split into the training and testing set. Therefore, the ALS dataset was assigned to the training set, allowing the machine learning algorithm to learn multiple disease expression signatures, which could further aid in differentiating AD from Non-AD subjects. Samples in the training set were up-sampled to prevent biasing the majority classes during model development. This resulted in “AD vs healthy” classification model consisting of 160 AD samples and 160 complimentary healthy control samples, and the “AD vs mixed controls” being trained on 6318 AD samples and 6318 non-AD samples. The “AD vs mixed controls” training set contains significantly more samples as the pooled controls consisted of 702 samples; therefore, the remaining 8 disease classes were up-sampled to the same sample size which totalled 6318 samples. The AD samples were then up-sampled to 6318 to balance the training set. An overview of subjects in the training and testing set is provided in Table 2.

**Table 2:** Overview Training and Testing set subjects

| Dataset | Training Set |  | Testing Set |
| --- | --- | --- | --- |
|  | AD vs healthy control | AD vs mixed control |  |
| Alzheimer's Disease | 160* | 6318 (160*) | 127 |
| Parkinson's Disease | 0 | 702 (50) | 40 |
| Multiple Sclerosis | 0 | 702 (122*) | 25 |
| Schizophrenia | 0 | 702 (97*) | 56* |
| Bipolar Disorder | 0 | 702 (28) | 20 |
| Cardiovascular Disease | 0 | 702 (235*) | 108 |
| Rheumatoid Arthritis | 0 | 702 (353) | 88* |
| Chronic Obstructive Pulmonary Disease | 0 | 702 (127) | 88 |
| ALS | 0 | 702 (52) | 0 |
| Pooled Controls | 160 (127*) | 702* | 262 |

Entire datasets from each disease were assigned to either the “Training Set” for classification model development or the “Testing Set” for validation purposes. Datasets with a larger number of diseased
subjects were prioritised into the training set to increase discovery. Two classification models were developed, the first was developed using only the 160 AD and associated 127 control samples, which were up-sampled to 160 to develop the “AD vs healthy control” classification model. The pooled controls in the “AD vs healthy control” training set originates only from AD datasets and can be regarded as cognitive healthy controls. The second classification model was developed using all datasets and diseases available in the training set and is referred to as the “AD vs mixed control” classification model, where samples in the minority classes are up-sampled to match the 702 samples in the pooled controls. The mixed control group totalled 6318 samples. Therefore, the AD group was up-sampled to 6318 to create a balanced training set. Sample numbers provided in brackets are before up-sampling. Sample numbers with an asterisk (\*) indicates multiple datasets were available, and subject numbers shown are a sum across these datasets.

### “AD vs healthy control” classification model development and performance

The “AD vs healthy control” classification model was developed using only the two AD datasets (GSE63060 and E-GEOD-6613) available in the training set, which after up-sampling consisted of 160 AD and 160 cognitive healthy controls. The model was initially built using default parameters which selected 126 predictive features from the available 1681 genes, resulting in a cross-validation test logloss mean of 0.47 (0.21 SD). Further refinement of the model identified 74 predictive genes and the optimum hyperparameters as **eta**=0.12, **max_depth**=10, **gamma**=0, **min_child_weight**=1, **subsample**=1, **colsample_bytree**=1, **alpha**=0, **lambda**= 1 and **nrounds** =63, which improved the test logloss mean to 0.27 (0.1 SD).

The “AD vs healthy control” classification model was validated on the independent testing set and achieved a sensitivity of 58.0%, specificity of 30.0% and a balanced accuracy of 44.3% (additional classification performance metrics are provided in Table 3). The probability predictions of individual samples in the testing set is illustrated in Figure 3a, where misclassification can be observed in all diseases and controls, demonstrating an increased false positive rate and the inability of the classification model to confidently assign a positive (0) or negative (1) class to each subject.

**Table 3:** Classification model performance

|  | “AD vs healthy control”<br>classification model | “AD vs mixed control”<br>classification model |
| --- | --- | --- |
| **Sensitivity** | 58.3% | 46.5% |
| **Specificity** | 30.3% | 95.6% |
| **Balanced Accuracy** | 44.3% | 71.0% |
| **PPV** | 13.4% | 66.3% |
| **NPV** | 79.7% | 90.6% |
| **AUC** | 0.49 | 0.84 |
| **AUC Rating** | Test not useful | Very good |
| **CUI +ve** | 0.08 | 0.31 |
| **CUI +ve Rating** | Poor | Poor |
| **CUI -ve** | 0.24 | 0.87 |
| **CUI -ve Rating** | Poor | Excellent |

**Figure 3:**
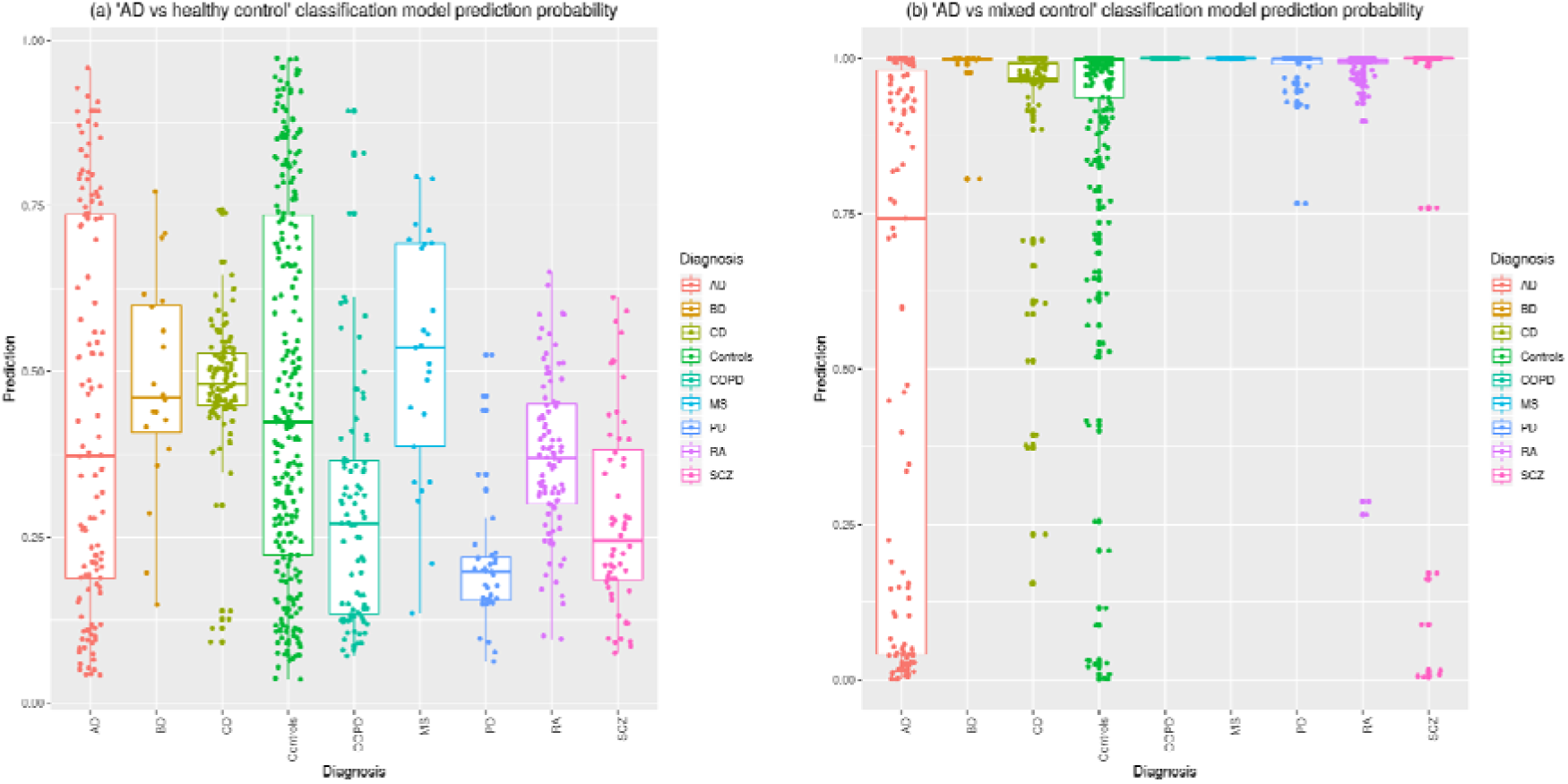
Illustrates the probability prediction of samples from the testing set being AD (0) or non-AD (1). Subjects in the testing set represent a heterogeneous ageing population with subjects being clinically diagnosed with various mental-health related disorders, neurodegenerative diseases, age-related diseases or are relatively healthy. a) illustrates the confidence of the “AD vs healthy control” classification model distinguishing subjects in the testing set and b) illustrates the confidence of the “AD vs mixed control” classification model in predicting the same testing set. Controls represent pooled non-diseased subjects from all datasets. Diseases are abbreviated as follows; AD = Alzheimer’s disease, BD = Bipolar disease, CD = Coronary Artery disease, COPD = Chronic Obstructive Pulmonary Disease, MS = Multiple Sclerosis, PD = Parkinson’s Disease, RA = Rheumatoid Arthritis and SCZ = Schizophrenia.

A ROC curve was generated for the “AD vs healthy control” classification model performance (Figure 4), which demonstrates a low TP rate in comparison to random and the AUC score of 0.49 suggests this “test is not useful” as a diagnostic test. The clinical utility values (CUI +ve = 0.08, CUI −ve = 0.24) mirrors the AUC score interpretation, as the CUI values suggest the classification model is “poor” at detecting the presence and absence of AD and based on current validation results, has no real clinical utility.

**Figure 4:**
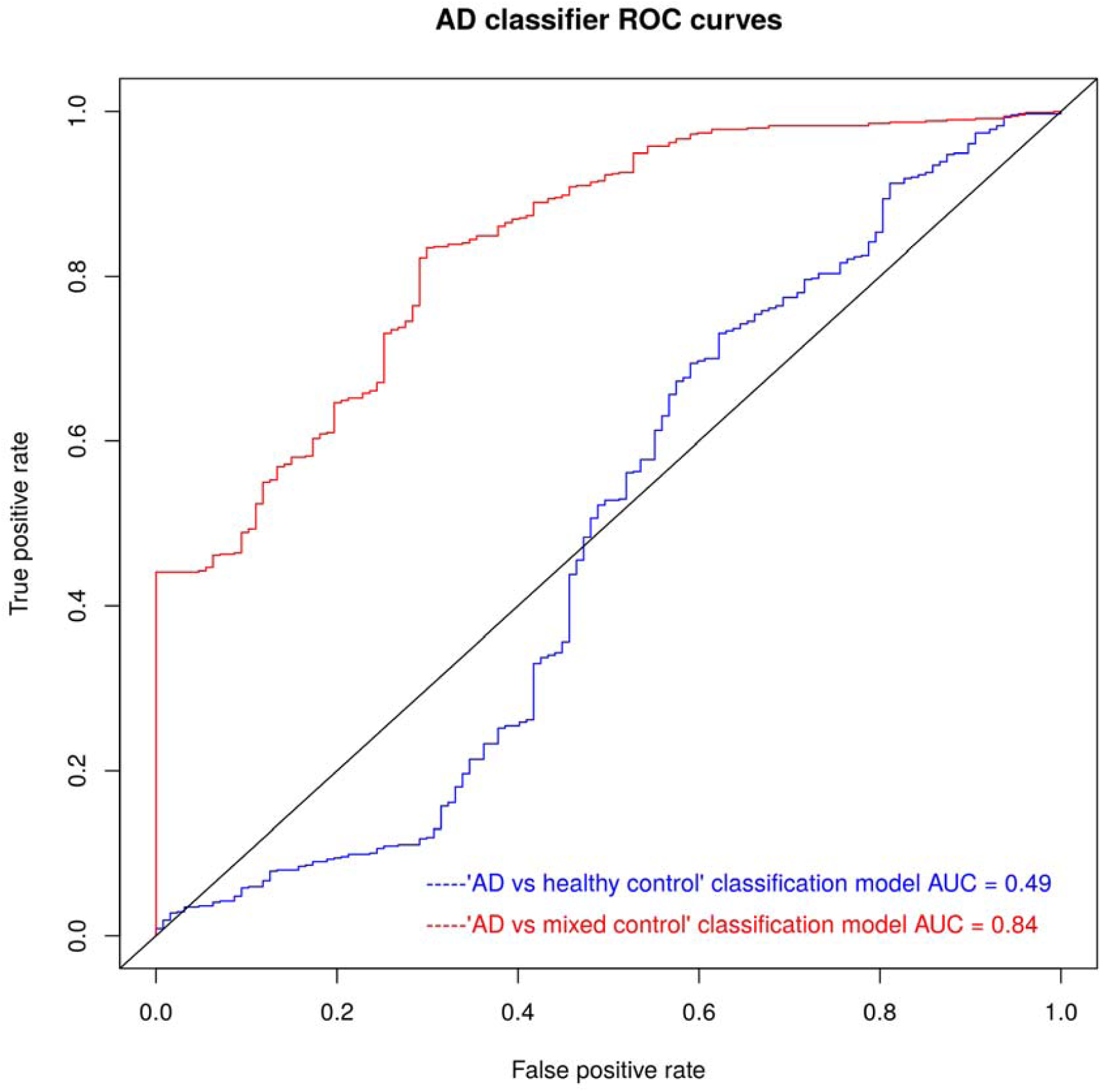
AD classification model ROC curves. The blue line represents the typical “AD vs healthy control” classification model’s ROC curve which is trained using AD and complimentary healthy control subjects. The red line represents the “AD vs mixed control” classification model’s ROC curve which is trained using AD and non-AD diseased and healthy control subjects. The ROC curves were generated from the performance of both classification models in the testing set and demonstrate an improved TP and FP rate with the “AD vs mixed control” classification model, which achieved an improved AUC value of 0.84 when compared to the typical “AD vs healthy” classification approach which achieved an AUC score of 0.49

### “AD vs mixed control” classification model development and performance

The “AD vs mixed control” classification model was developed on the entire training set, which after up-sampling consisted of 6318 AD and 6318 non-AD subjects. The classification model was built using default parameters which selected 231 genes from the available 1681 genes as predictive features and resulted in cross-validation test logloss mean of 0.015 (0.009 SD).). Further refinement of the model identified 28 predictive features, **eta**=0.08, **max_depth**=6, **gamma**=0, **min_child_weight**=1, **subsample**=1, **colsample_bytree**=1, **alpha**=0, **lambda**=0.9 and **nrounds**=139 as the optimum parameters which improved the cross-validation test logloss mean to 0.009 (0.005 SD).

The “AD vs mixed control” classification model was further validated on the testing set and achieving 46.5% sensitivity, 95.6% specificity, and a balanced accuracy of 71.0% (additional classification performance metric are provided in Table 3). The performance of this classification model improves on the typical “AD vs healthy control” classification model in all performance metrics, except for sensitivity, where a decrease in performance is observed from 58% to 46.5%. Nevertheless, due to the “AD vs mixed control” classification model predicting less false positives, an increase in PPV (66.3%) is observed when compared to the “AD vs healthy control” classification model (PPV = 13.4%). Furthermore, as illustrated in Figure 3b, the probability predictions for individuals in the testing set are more correctly and confidently predicted when compared to the typical “AD vs healthy control” classification model, only misclassifying 21 pooled controls (8% of total pooled controls), 2 CD (2% of CD subjects), and 7 SCZ (13% of SCZ subjects) as AD. The “AD vs mixed control” classification model ROC curve (Figure 4) achieves an improved AUC score of 0.84 which translates to a “very good” diagnostic test, however, the clinical utility values (CUI +ve = 0.31 and CUI −ve = 0.87) suggests this classification model is “poor” in the detection of AD but “excellent” to rule out “AD”.

### “AD vs mixed control” classification model’s predictive features

The variable importance was calculated for the 28 predictive genes (gene list provided in Supplementary Table 1) with the 20 most predictive genes illustrated in Figure 5. The gene **LDHB** provides the greatest predictive value with a relative importance value of 0.7. GSEA performed on the 28 genes identified four biological processes significantly enriched; **WNT ligand biogenesis and trafficking** (p-value= 9.56e-4, q-value=0.04), **Alzheimer disease** (p-value= 3.70e-3, q-value=0.05), **Herpes simplex infection** (p-value= 4.61e-3, q-value=0.05), and **Huntington disease** (p-value= 5.19e-3, q-value=0.05). Additional information on gene overlap between the 28 predictive genes and biological pathways is provided in Supplementary Table 2.

**Figure 5:**
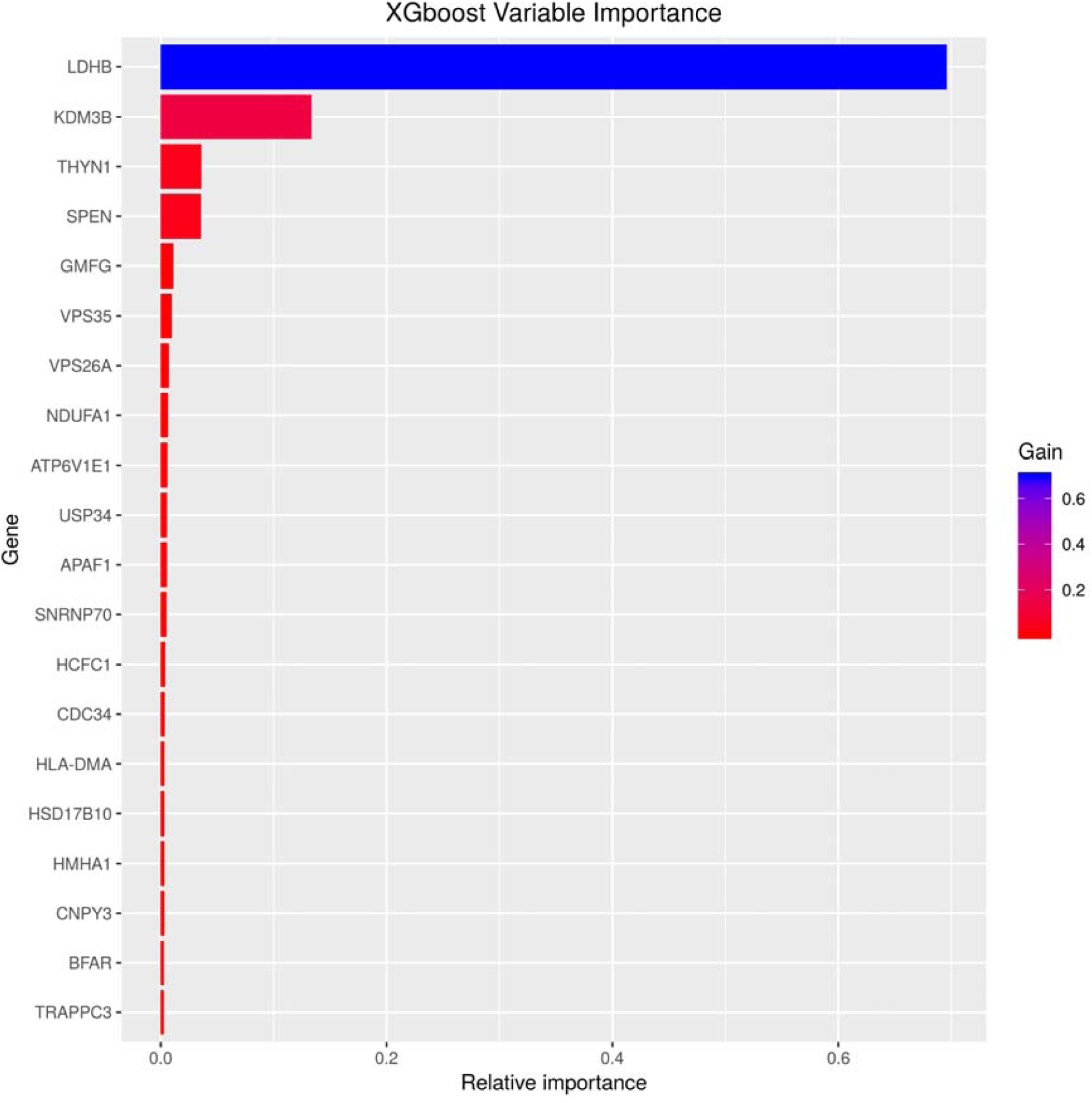
Relative Importance of the 20 most predictive genes for the “AD vs mixed control” classification model

## Discussion

Previous attempts to identify blood-derived gene expression profiling for AD diagnosis have relied on the typical approach of training machine learning algorithms on AD and cognitively healthy subjects only, inadvertently leading to classification models learning expression signatures that may be of general illness rather than being disease-specific. Validating such a classification model in a heterogeneous ageing population may fail to distinguish AD from similar mental health disorders, neurodegenerative diseases, age-related disorders and cognitive healthy individuals. To address this issue, this study developed an “AD vs mixed control” classification model based on a training set comprised of AD, PD, MS, BD, SCZ, CD, RA, COPD, ALS and a set of pooled healthy individuals totalling 1926 subjects. The individual classes within the mixed control group varied in sample size, with the pooled controls representing the largest class consisting of 702 samples. Therefore, to avoid sampling bias during the classification model development, the individual classes in the mixed control group were each up-sampled with replacement to 702, which totalled 6318 samples. The AD group were then up-sampled to 6318, classification model developed, optimised and evaluated in an external independent cohort comprised of similar diseases and controls totalling 814 subjects.

The typical approach of developing a classification model trained on AD and complimentary cognitive healthy control subjects produced a model with a sensitivity of 58.3% in an independent cohort of 127 AD subjects. The performance of this model is slightly better than a previous attempt which attained a sensitivity of 56.8% when validated in an independent testing set of 118 AD subjects [10]. However, when evaluating this typical AD classification model in a heterogeneous ageing population, a process often neglected in previous studies, very low specificity of 30.3% was attained which equated to a low PPV of only 13.4%. PD is the second most common cause of dementia [1] and was seen to be most misclassified as AD. However, since misclassification was observed in all groups including large portions of the controls, this classification model is most likely not capturing signals of AD, dementia or general illness, but is most likely a result of technical noise, individual study batch effects and overfitting. This is mirrored in the model’s performance metrics which translates to a “poor” clinical utility in detecting the presence and absence of AD. Overall, the typical approach of AD classification model development failed to accurately distinguish AD subjects in a heterogeneous ageing population consisting of PD, MS, BD, SCZ, CD, RA, COPD, ALS and relatively healthy controls.

In contrast, the “AD vs mixed control” classification model attained a validation PPV of 66.3% and NPV of 90.6% on the same testing set, which outperforms the validation PPV of 13.4% and NPV of 79.9% achieved by the “AD vs healthy control” classification model. However, this improvement was at the cost of sensitivity, which was reduced from 58.3% (“AD vs healthy control”) to 46.4% (“AD vs mixed control”). Nevertheless, an overall increase in the clinical utility of the “AD vs mixed control” classification model was measured and according to the recommended CUI interpretations in [24], the model is “poor” in “ruling in” AD but “excellent” in “ruling out” AD.

The performance of the “AD vs mixed control” classification model can be suggested to be superior due to the increased number of samples in the training set. The “AD vs healthy control” classification model was developed using 160 AD samples while the “AD vs mixed controls” classification model was developed using 6318 AD samples. However, it is important to note the AD samples in both training sets originated from the same subjects, with AD sample numbers in the “AD vs mixed control” training set up-sampled to account for the variation of sample sizes across the individual classes in the mixed control group. Therefore, the increased performance achieved by the “AD vs mixed control” classification model is most likely the result of incorporating additional related neurological and age-related disorders into the classification model development process, which aided in the identification of a more AD-specific expression signature than the typical approach of using only AD and corresponding control samples. Although this improved the ability to distinguish AD from other related diseases and cognitively healthy controls, the sensitivity of the model was reduced and needs to be further enhanced for this type of research to be beneficial in the clinical setting.

The underlying replication of predictive genes across blood-based transcriptomic biomarker studies are inconsistent [44]–[47]. Nevertheless, sets of genes within independent studies have been able to consistently distinguish AD from complimentary controls [44]. Therefore, the predictive features in this study warrant further investigation to assess their biological relevance to AD. The “AD vs mixed control” classification model differentiates AD from other diseases and healthy controls using the relationship of 28 genes. GSEA identified “**Herpes simplex infection**” as one of the biological pathways being significantly enriched prior to multiple corrections, with an overlap of 3 genes (**CDC34, HCFC1 and HLA-DMA**). This suggests gene expression changes associated with this process are measurable in blood and may contribute towards identifying AD subjects. Pathogenic viral components have been long suspected of playing an essential role in the onset and progression of AD. A recent study identified common viral species in normal and ageing brains, with an increased human herpesvirus 6A and human herpesvirus 7 in AD brains [48]. In addition, a recent transcriptomic meta-analysis study identified genes involved with “**interspecies interactions**” were specifically enriched in AD brains when taking into account expression changes in related neurological disorders [49]. The current observation extends the previous suggestions of viral involvement in AD brains to blood, as the expression of blood-derived genes involved in the “**Herpes simplex infection”** can be used to distinguish AD from other neurological diseases and control subjects in this study.

Age is one of the most significant risk factors for AD, and the prevalence of the disease is known to increase with age. A meta-analysis study investigating blood transcriptional changes associated with age in 14,983 humans, identified 1,496 differentially expressed genes with chronical age [50], of which three genes (**LDHB, AARS** and **ABR**) are in the “AD vs mixed control” classification model’s 28 predictive genes. The classification models most predictive gene **LDHB** is ranked 28^th^ in the meta-analysis study and was also observed to be negatively associated with age in the brain, specifically the frontal cortex and cerebellum [50]. The datasets used in this study were publicly available, and as such, were accompanied with limited phenotypic information, including age. Therefore, age was not accounted for during the classification model developmental process. However, as this study uses a variety of age-related diseases, in addition to the 3 AD datasets, and study designs generally incorporate complementary age-matched controls, it is highly unlikely the classification model is predicting age alone but is more likely using a combination of signals including age to distinguish AD. Without age information for all subjects, this study is unable to conclude how age is influencing the model prediction process.

All data used in this study were publicly available, and as such, many were accompanied by limited phenotypic information, including basic sex information, which was predicted based on gene expression when missing. Therefore, this study was unable to incorporate additional phenotypic information during the classification model building process, which has been shown to improve model performance [10]. Information such as comorbidities, age and medications are unknowns which could be affecting performance in this study. For instance, control subjects in this study that originated from non-AD datasets were screened negative for their corresponding disease of interest but were not screened for cognitive function. i.e. control subjects from the CD datasets were included in their retrospective dataset if they did not have CD, they were not necessarily checked for cognitive impairment. Therefore, some misclassified control subjects may indeed be on the AD spectrum, and it’s important to note subjects from the pooled control group were most misclassified as AD by the “AD s mixed control” classification model. However, it is also important to note the training set used to develop the “AD vs mixed control” classification model also contains these controls which have not been screened for AD. If these controls or age-related disease subjects are comorbid with AD, the classification model may have inadvertently learned to be biased towards a subgroup of AD subjects with no comorbid with any other disease, hence the low sensitivity validation performance when introducing additional datasets into the classification model developmental process.

This study involved a number of subjects clinically diagnosed with age-related diseases, and most likely, are on some sort of therapeutic treatment to manage or treat the underlying disease, another piece of vital information generally missing from publicly available datasets and from this study. As therapeutic drugs have been well-known to affect gene expression profiling, including memantine, a common drug used to treat AD symptoms [51], the “AD vs mixed control” classification model may have inadvertently learnt gene expression perturbations due to therapeutic treatment rather than disease biology, and would, therefore, fail in the clinical setting to diagnose AD subjects who are not already on medication. To address this issue along with co-morbidity, clear and detailed phenotypic information would be needed for all subjects, which is encouraged for future studies planning to submit genetic data to the public domain.

This study used datasets generated on 11 different microarray BeadArrays, resulting in datasets ranging from 22277-54715 probes prior to any QC. Coupled with differences in BeadArrays designs across platforms, the overlap of genes was drastically reduced to 1681 common “reliable detected” genes across all datasets. This ensured gender-specific expression changes were captured; however, this may have also inadvertently lost some disease-specific changes. To address this issue, these subjects need to be expression profiled on the same microarray platform and ideally the same expression BeadArray, which currently doesn’t exist in the public domain. The advances in sequencing technologies which can capture expression changes across the whole transcriptome can potentially solve this issue and future studies are encouraged to replicate this study design with RNA-Seq data with detailed phenotypic information when/if available, albeit, this may bring new challenges.

## Conclusion

This study relied on publicly available microarray gene expression data, which too often lacks detailed phenotypic information for appropriate data analysis and needs to be addressed by future studies. Nevertheless, with the available phenotypic information and limited common “reliably detected” genes across the different microarray platforms and BeadArrays, this study demonstrated the typical approach of developing an AD blood-based gene expression classification model using only AD and complimentary healthy controls fails to accurately distinguish AD from a heterogeneous ageing population. However, by incorporating additional related neurological and age-related diseases into the classification model development process can result in a model with improved “predictive power” in distinguishing AD from a heterogeneous ageing population. Nevertheless, further improvement is still required in order to identify a robust blood transcriptomic signature more specific to AD.

## Author contributions statement

SJN and RJBD proposed the initial concept of the study. HP, SJN and RJBD developed the detailed study design, with support from RI and DS for machine learning aspect. HP acquired and analysed data. All authors contributed to interpretation of results. HP, RJBD and SJN drafted the manuscript.

## Acknowledgements

This study presents independent research supported by the NIHR BioResource Centre Maudsley at South London and Maudsley NHS Foundation Trust (SLaM) & Institute of Psychiatry, Psychology and Neuroscience (IoPPN), King’s College London. The views expressed are those of the author(s) and not necessarily those of the NHS, NIHR, Department of Health or King’s College London.

RJBD and SJN are supported by 1. Health Data Research UK, which is funded by the UK Medical Research Council, Engineering and Physical Sciences Research Council, Economic and Social Research Council, Department of Health and Social Care (England), Chief Scientist Office of the Scottish Government Health and Social Care Directorates, Health and Social Care Research and Development Division (Welsh Government), Public Health Agency (Northern Ireland), British Heart Foundation and Wellcome Trust. 2. The National Institute for Health Research University College London Hospitals Biomedical Research Centre.

